# Candidate gene scan for Single Nucleotide Polymorphisms involved in the determination of normal variability in human craniofacial morphology

**DOI:** 10.1101/060814

**Authors:** Mark Barash, Philipp E. Bayer, Angela van Daal

**Author notes:** Corresponding author Corresponding author postal address: Centre for Forensic Sciences, School of Mathematical and Physical Sciences, Faculty of Science University of Technology Sydney, Building 4, Thomas St, Broadway NSW 2007, Australia. Emails: Mark Barash; Philipp E. Bayer; Angela van Daal.

## Abstract

Despite intensive research on genetics of the craniofacial morphology using animal models and human craniofacial syndromes, the genetic variation that underpins normal human facial appearance is still largely elusive. Recent development of novel digital methods for capturing the complexity of craniofacial morphology in conjunction with high-throughput genotyping methods, show great promise for unravelling the genetic basis of such a complex trait.

As a part of our efforts on detecting genomic variants affecting normal craniofacial appearance, we have implemented a candidate gene approach by selecting 1,201 single nucleotide polymorphisms (SNPs) and 4,732 tag SNPs in over 170 candidate genes and intergenic regions. We used 3-dimentional (3D) facial scans and direct cranial measurements of 587 volunteers to calculate 104 craniofacial phenotypes. Following genotyping by massively parallel sequencing, genetic associations between 2,332 genetic markers and 104 craniofacial phenotypes were tested.

An application of a Bonferroni–corrected genome–wide significance threshold produced significant associations between five craniofacial traits and six SNPs. Specifically, associations of nasal width with rs8035124 (15q26.1), cephalic index with rs16830498 (2q23.3), nasal index with rs37369 (5q13.2), transverse nasal prominence angle with rs59037879 (10p11.23) and rs10512572 (17q24.3), and principal component explaining 73.3% of all the craniofacial phenotypes, with rs37369 (5p13.2) and rs390345 (14q31.3) were observed.

Due to over-conservative nature of the Bonferroni correction, we also report all the associations that reached the traditional genome-wide p-value threshold (<5.00E-08) as suggestive. Based on the genome-wide threshold, 8 craniofacial phenotypes demonstrated significant associations with 34 intergenic and extragenic SNPs. The majority of associations are novel, except *PAX3* and *COL11A1* genes, which were previously reported to affect normal craniofacial variation.

This study identified the largest number of genetic variants associated with normal variation of craniofacial morphology to date by using a candidate gene approach, including confirmation of the two previously reported genes. These results enhance our understanding of the genetics that determines normal variation in craniofacial morphology and will be of particular value in medical and forensic fields.

**Author Summary:** There is a remarkable variety of human facial appearances, almost exclusively the result of genetic differences, as exemplified by the striking resemblance of identical twins. However, the genes and specific genetic variants that affect the size and shape of the cranium and the soft facial tissue features are largely unknown. Numerous studies on animal models and human craniofacial disorders have identified a large number of genes, which may regulate normal craniofacial embryonic development.

In this study we implemented a targeted candidate gene approach to select more than 1,200 polymorphisms in over 170 genes that are likely to be involved in craniofacial development and morphology. These markers were genotyped in 587 DNA samples using massively parallel sequencing and analysed for association with 104 traits generated from 3-dimensional facial images and direct craniofacial measurements. Genetic associations (p-values<5.00E-08) were observed between 8 craniofacial traits and 34 single nucleotide polymorphisms (SNPs), including two previously described genes and 26 novel candidate genes and intergenic regions. This comprehensive candidate gene study has uncovered the largest number of novel genetic variants affecting normal facial appearance to date. These results will appreciably extend our understanding of the normal and abnormal embryonic development and impact our ability to predict the appearance of an individual from a DNA sample in forensic criminal investigations and missing person cases.

## Introduction

The human face is probably the most commonly used descriptor of a person and has an extraordinary role in human evolution, social interactions, clinical applications as well as forensic investigations. The influence of genes on facial appearance can be seen in the striking resemblance of monozygotic twins as well as amongst first degree relatives, indicating a high heritability [1, 2].

Uncovering the genetic background for regulation of craniofacial morphology is not a trivial task. Human craniofacial development is a complex multistep process, involving numerous signalling cascades of factors that control neural crest development, followed by a number of epithelial-mesenchymal interactions that control outgrowth, patterning and skeletal differentiation, as reviewed by Sperber et. al. [2]. The mechanisms involved in this process include various gene expression and protein translation patterns, which regulate cell migration, positioning and selective apoptosis, subsequently leading to development of specific facial prominences. These events are precisely timed and are under hormonal and metabolic control. Most facial features of the human embryo are recognizable from as early as 6 weeks post conception, developing rapidly *in utero* and continuing to develop during childhood and adolescence [3, 4]. Development of the face and brain are interconnected and occur at the same time as limb formation. Facial malformations therefore, frequently occur with brain and limb abnormalities and vice versa. Genetic regulation of craniofacial development involves several key morphogenic factors such as *HOX*, *WNT*, *BMP*, *FGF* as well as hundreds of other genes and intergenic regulatory regions, incorporating numerous polymorphisms [2]. The SNPs involved in craniofacial diseases may in fact influence the extraordinary variety of human facial appearances, in the same way that genes responsible for albinism have been shown to be involved in normal pigmentation phenotypes [5]. Additionally, non-genetic components such as nutrition, climate and socio-economic environment may also affect human facial morphology via epigenetic regulation of transcription, translation and other cellular mechanics. To date, both the genetic and even more so, the epigenetic regulation of craniofacial morphology shaping are poorly understood.

The genetic basis of craniofacial morphogenesis has been explored in numerous animal models with multiple loci shown to be involved [2]. The majority of human studies in this field have focused on the genetics of various craniofacial disorders such as craniosynostosis and cleft lip/palate [6, 7], which may provide a link to regulation of normal variation of the craniofacial phenotype, as for example observed between cleft-affected offspring and the increase of facial width seen in non-affected parents [8]. These studies have identified several genes with numerous genetic variants that may contribute to normal variation of different facial features, such as cephalic index, bizygomatic distance and nasal area measurements [9–11]. Studies of other congenital disorders involving manifestation of craniofacial abnormalities such as Alagille syndrome (*JAG1* and *NOTCH2* gene mutations), Down syndrome (chromosome 21 trisomy - multiple genes), Floating-Harbor syndrome (*SRCAP* gene mutations) and Noonan syndrome (mutations in various genes such as *PTPN11* and *RAF1)* provide additional information on the candidate genes potentially involved in normal craniofacial development [12–17].

In recent years, new digital technologies such as 3-Dimentional laser imaging have been used in numerous anthropometric studies. 3-D laser imaging allows accurate and rapid capture of facial morphology, providing a better alternative to traditional manual measurements of craniofacial distances [18–20]. The high-throughput genotyping technologies and digital methods for capturing facial morphology have been used in a number of recent studies that demonstrated a link between normal facial variation and specific genetic polymorphisms [21–23]. Despite these promising results, our current knowledge of craniofacial genetics is sparse.

This study aims to further define the polymorphisms associated with normal facial variation using a candidate gene approach. The advantage of a candidate gene approach over previous genome wide association studies (GWAS) is that it focuses on genes, which have previously been associated with craniofacial embryogenesis or inherited craniofacial syndromes, rather than screening hundreds of thousands of non-specific markers. This approach aims to increase the chances of finding significant associations between SNPs and visible traits and requires fewer samples for robust association analysis [24, 25].

In the current study, 32 anthropometric landmarks were recorded from 3-D facial scans of 587 volunteers from general Australian population (Gold Coast, Queensland). Additionally, three direct cranial measurements using a calliper were made and two facial traits (ear lobe and eye lid morphology) were recorded. Both the direct measurements and the Cartesian coordinates of the anthropometric landmarks were used to calculate 92 craniofacial distances. The calculation of 10 principal components based on the craniofacial measurements was performed in order to obtain a more simplified representation of the facial shape. The associations between 104 of the total craniofacial traits and 2,332 genetic markers were tested.

This research aims to assist in uncovering the genetic basis of normal craniofacial morphology variation and will enhance our understanding of craniofacial embryogenetics. These findings could be useful in building models to predict facial appearance from a forensic DNA sample where no suspect has been identified, thereby providing valuable investigative leads. It could also assist in identifying skeletal remains by allowing more accurate facial reconstructions.

## Results

### 3D measurements precision study

In the last decade 3D scanning systems have been extensively used in anthropometric studies as well as in medical research [18, 20, 64, 65]. The Minolta Vivid V910 3D scanner has been demonstrated to have accuracy to a level of 1.9 ± 0.8 mm [66] and 0.56 ± 0.25 mm [67], making it suitable for the present study since it should provide an accurate representation of facial morphology. However, the allocation of anthropometric facial landmarks can be challenging, especially when tissue palpating is not possible. Reproducibility of the landmark precision was assessed on fifteen 3D facial images through assessment of 85 facial measurements, including linear and angular distances and ratios between the linear distances at two separate times. The period between the analyses varied from one to six months. The mean difference (MD) was calculated as the discrepancy between the first and the second measurement. The measurement error (ME) was calculated as the standard deviation of the MD divided by square root of 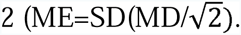

In general, the nasal area distances, which involved nasion, pronasale, subnasale and alare landmarks showed greater reproducibility, while the measurements involving paired landmarks, such as gonion and zygion demonstrated higher variance. This result can be explained by easier allocation of nasal area landmarks, compared with gonion and zygion [29]. Overall the median difference (MD) between two measurements for linear distances in 15 images ranged between 0.76 mm (ME ±0.27) and 2.80 mm (ME ±0.99); for angular distances between 0.38 mm (ME ±0.96) and 3.75 mm (ME ±0.40) and for facial indices (ratios) between 0.46 mm (ME ±1.08) and 2.98 mm (ME ±1.95) respectively. The lower reproducibility in the angular distances and indices can be explained by a higher number of landmarks (hence variability in allocation of x, y and z coordinates) needed for their calculation (three and four landmarks respectively). Nevertheless, our findings are concordant with the published results, which observed variance of 0.19 mm to 3.49 mm with a ME range of 0.55 mm to 3.34 mm for each landmark [19, 68].

### Candidate genes search and sequencing data quality control

The search for candidate genes and SNPs potentially involved in influencing normal craniofacial morphology variation initially focused on searching for genes involved in normal or abnormal craniofacial variation in humans and model organisms (Supplemental Table S1). As a complementary approach, a search for genetic markers with high Fst values (≥0.45) was implemented, based on the rationale that genes involved in craniofacial morphology regulation are likely to display significant differences in allele frequencies across populations.

The first approach has mainly focused on the Mouse Genome Informatics (MGI) database search using the keyword ‘craniofacial mutants’ and additional resources such as Online Mendelian Inheritance in Man (OMIM), GeneCards and AmiGO, using the keywords such as “craniofacial”, “craniofacial mutants”, “craniofacial anomalies”, “craniofacial dimorphism” and “facial morphology” (a detailed list of used resources is summarized in Supplemental Appendix S1). This search revealed a list of 2,891 genotypes and 7,956 annotations. A search of the ‘abnormal facial morphology’ sub-category resulted in 1,492 genotypes and 2,889 annotations. The final search of the ‘abnormal nose morphology’ of the previous sub-category revealed 219 genotypes with 310 annotations, representing approximately 150 genes.

In parallel, a search for high Fst markers, using previously published AIMs and web tools, such as ENGINES, resulted in identification of additional targets, for a total of 1,088 genes and intergenic regions (a detailed list of used resources is summarized in Supplemental Appendix S1).

However, manual examination revealed that 592 of these genes showed no apparent link with normal craniofacial development or malformations and were therefore excluded. The remaining 496 regions were further screened for non-synonymous and potentially functional SNPs, as well as SNPs with high population differentiation, which resulted in the shortlist of 269 genes and intergenic regions.

Subsequent analysis of these 269 genes/regions for functional annotation using the AmiGO Gene Onthology server [57], resulted in 177 candidate genes/regions, possessing 1,319 genetic markers involved in various stages of human embryonic development, including: embryonic morphogenesis, sensory organ development, tissue development, pattern specification process, tissue morphogenesis, ear development, tube morphogenesis, epithelium development, chordate embryonic development and morphogenesis of an epithelium (Supplemental Appendix S1). Notably, the majority of these markers are located in introns and intergenic regions.

In terms of molecular function, AmiGO showed that craniofacial candidate markers might be involved in a range of regulatory activities including: protein dimerization activity, chromatin binding, regulatory region DNA binding, sequence-specific DNA binding RNA polymerase II transcription factor activity, sequence-specific distal enhancer binding activity, heparin binding, RNA polymerase II core promoter proximal region sequence-specific DNA binding transcription factor activity involved in positive regulation of transcription, BMP receptor binding and transmembrane receptor protein serine/threonine kinase binding (Supplemental Appendix S1).

Subsequent analysis of candidate SNPs for mouse phenotype associations confirmed that orthologous candidate markers were previously detected in mouse models displaying abnormal morphology of the skeleton, head, viscerocranium and facial area, as well as specific malformations of the eye, ear, jaw, palate, limbs, digits and tail (data not shown).

In additional to craniofacial candidate SNPs, 522 markers, previously shown to be associated with pigmentation traits, such as eye, skin and hair colour were selected from the relevant literature. These markers were used to validate the results of the genetic association analyses of craniofacial traits.

The final candidate marker list was analysed using the GREAT platform to visualize the genomic context of amplicons covering targeted SNPs [69]. The analysis revealed that almost 99% of the genomic regions (which may cover multiple markers) are associated with one or two genes with approximately 62% of genomic regions located 0-500 kb downstream of a transcription start site (data not shown).

Targeted massively parallel sequencing of the 587 samples resulted in 9,051 genetic markers, with the majority of markers (>5,000) represented by rare polymorphisms of ≤1% minor allele frequency (MAF) (data not shown). The difference between the initial hot-spot SNP panel of candidate markers (n=6,945) and the actual sequencing output (n=9,051) was a result of identification of potentially novel and rare markers in individual DNA samples. Three of the 587 samples, did not produce high quality genotypes because of poor DNA quality or unsuccessful library and template preparation.

The SNPs were filtered by sequencing quality and by MAF. Data quality control was performed by removing markers of low genotype quality (GQ>10) and sequencing depth (DP>10X), which resulted in 8,518 markers (Supplemental Appendix S2). Further filtering of markers using a 2% MAF cut-off resulted in 3,075 markers (Supplemental Appendix S2). The decision to apply a slightly more stringent MAF threshold (2%) was made because of the sample size (n=587) and to reduce potential bias from rare SNPs (1% MAF). Since this may reduce the power of analysis, we analysed and compared both datasets and did not observed any significant difference. Additional filtering based on the HWE threshold of p-value ≥0.01 resulted in 2,332 markers. The mean sequencing depth for significantly associated markers in this study was 58 fold (±48.9 SD).

### Genetic association study

The association analyses were performed using a linear regression model, incorporating EIGENSTRAT-generated PCA as well as sex and BMI as covariates. The use of covariates in the statistical analysis aimed to reduce the risk of introducing confounding effects, which can result in false positive associations. While sexual dimorphism in the craniofacial morphology is well-known [70], BMI will also likely affect certain craniofacial traits, since the soft facial tissue may change significantly with weight gain or loss. Despite that, this potential confounding factor has to date been disregarded in association studies of normal craniofacial morphology. Age was not considered a significant covariate, given that average age of the subjects in this study was 27 (±8.9 SD). Nevertheless, the potential effect of age as a cofactor was assessed on three craniofacial traits and found to be not significant (data not shown).

While the majority of current GWA studies rely on a p-value <5.00E-08 significance threshold, some publications suggest this threshold may be too stringent, especially for complex traits that are regulated by a large number of small effect alleles [75, 76]. In contrast to GWAS, candidate gene studies undertake a more focused genetic strategy, concentrating on a relatively limited number of putative markers. As this study analysed a significantly lower number of SNPs than usual GWA-studies, we could use a higher p-value cut-off since the smaller sample size means the probability of false positive at extremely low p-values is itself lower. Nevertheless, we decided to keep the traditional GWAS p-value significance threshold (<5.00E-08) in order to reduce the possibility of detecting false positive results.

In addition, we subsequently applied a more stringent Bonferroni – corrected threshold in order to minimize the chance of detecting spurious associations. Following the association analysis of 104 craniofacial phenotypes with 2,332 genetic markers, the significance threshold based on the Bonferroni correction with a desired α of 0.05 would be 2.06E-07 (=0.05/(2,332*104)).

However, it should be emphasized that the Bonferroni correction is widely considered over-conservative, especially in the case of complex phenotypic traits with small individual effects of each allele. Considering that our results confirm the previously published findings, we believe the GWAS p-value threshold is conservative enough to avoid or at least significantly reduce potentially spurious associations. Following this rationale, we report all the variants, which met the unadjusted genome-wide association p-value threshold as suggestive. We believe these findings are useful for the future studies focusing on genetics of normal craniofacial morphology.

The results of the association analyses of the craniofacial traits are summarized in Table 1 and Supplemental Figs. S1-S16. In general, following the application of a stringent Bonferroni-corrected GWAS threshold (adjusted p-value <1.6E-07), we observed five craniofacial traits being associated with six genomic markers. Specifically, nasal width with rs8035124 (p-value 1.74E-07, Beta=1.366, SE=0.209), cephalic index with rs16830498 (p-value 8.67E-08, Beta=3.005, SE=0.4518), nasal index with rs37369 (p-value 1.43E-07, Beta=4.025, SE=0.6124), transverse nasal prominence angle with rs59037879 (p-value 6.07E-09, Beta=4.765, SE=0.6685) and rs10512572 (p-value 1.57E-08, Beta=1.505, SE=0.2171), and principal component (EV=1391.99) with rs37369 (p-value 2.85E-08, Beta=-0.021, SE=0.003079) and rs390345 (p-value 8.55E-08, Beta=-0.0184, SE=0.002768). The polymorphisms: rs16830498, rs59037879 and rs390345 are intronic variants in *CACNB4*, *ZEB1* and *FOXN3* respectively; rs37369 is a missense mutation in the *AGXT2* gene and rs8035124 and rs10512572 are intergenic variants in 15q12.2 and 17q21.33 chromosomal locations respectively.

**Table 1.**
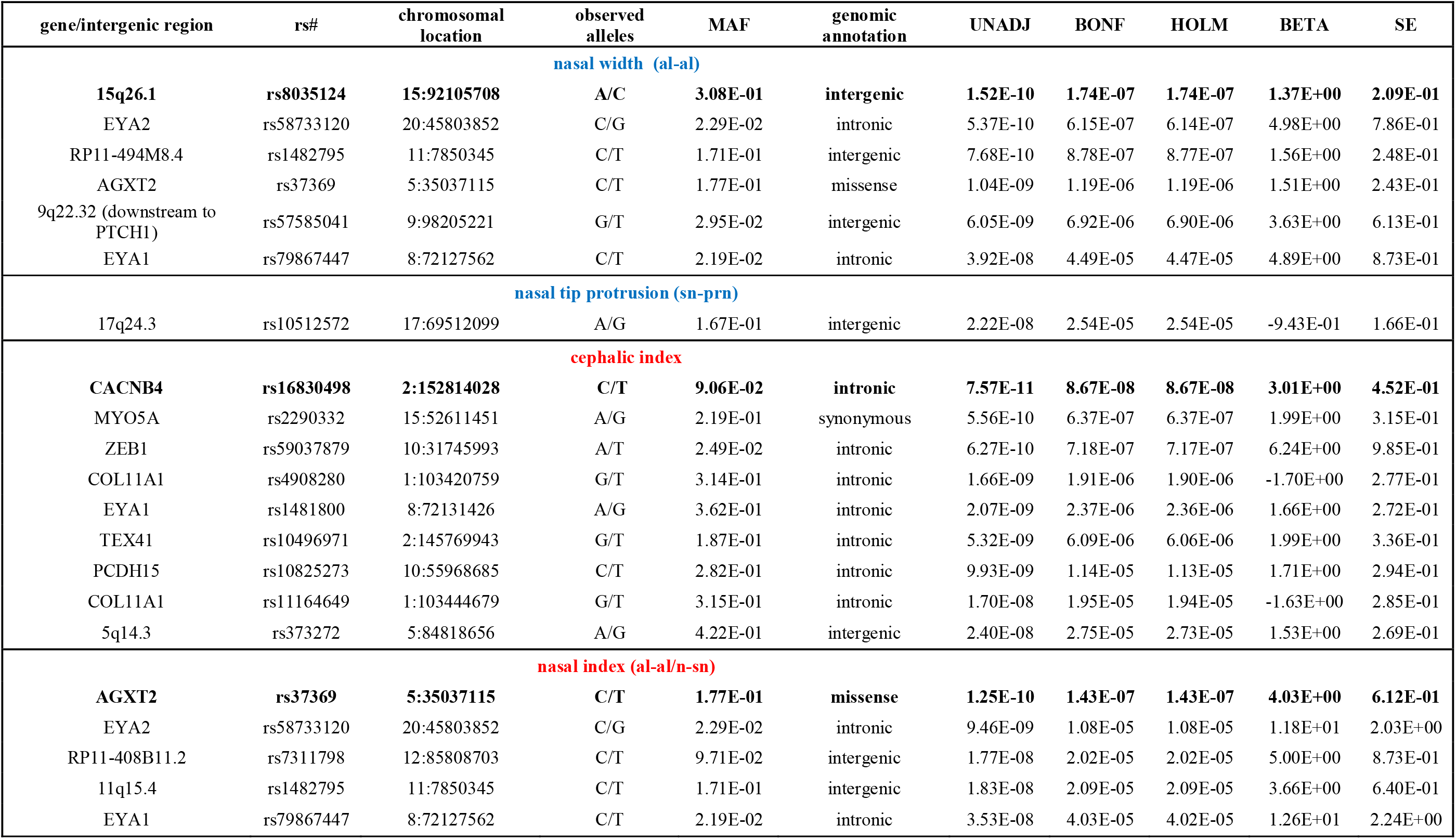
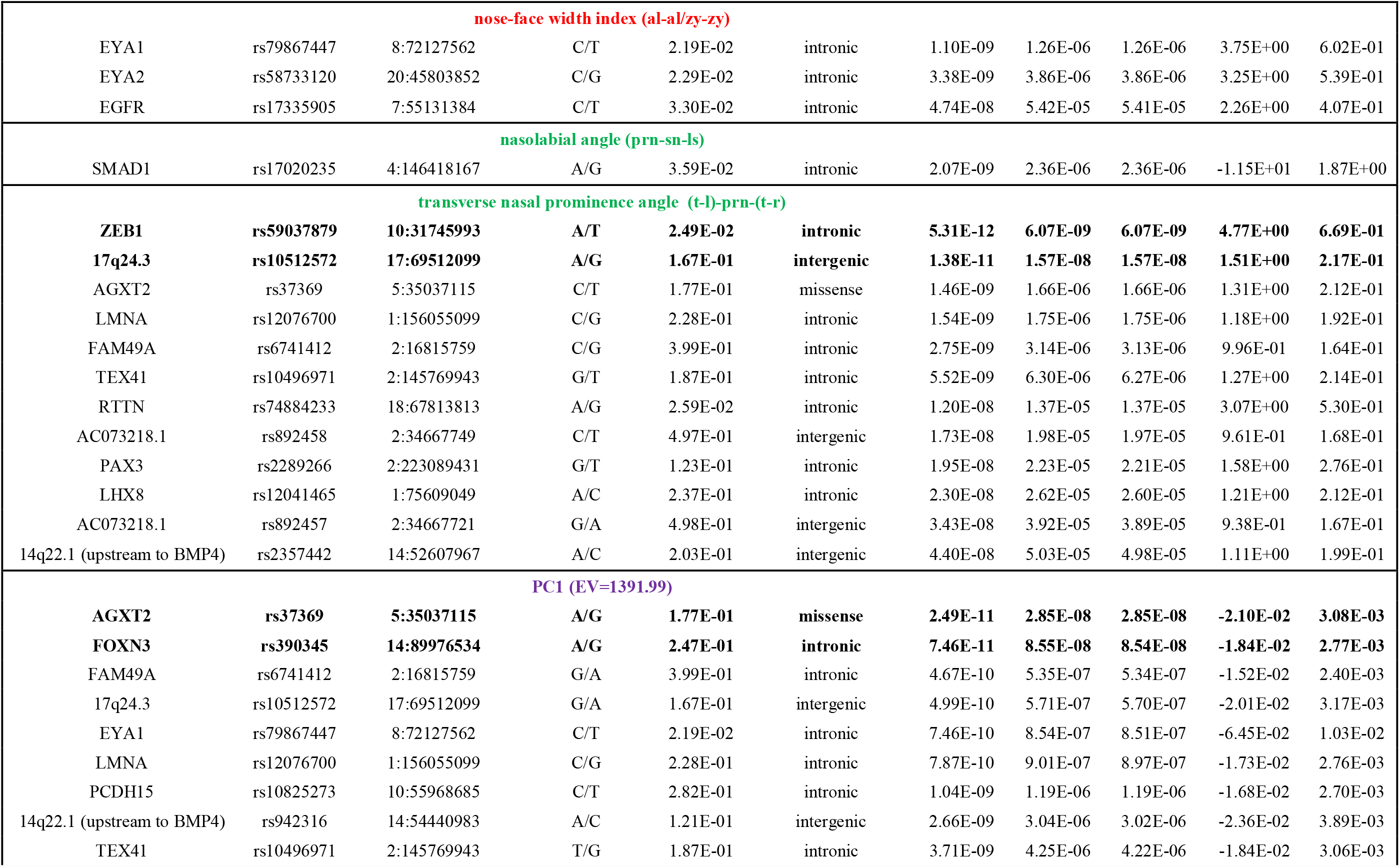
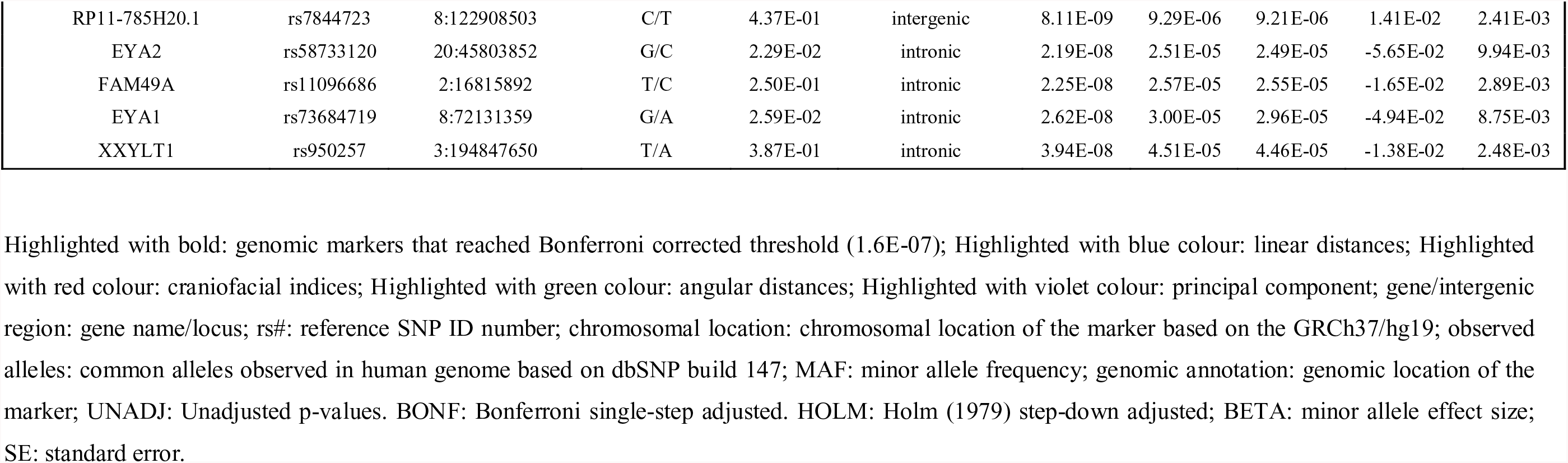
Results of genetic association analyses between candidate SNPs and craniofacial traits, including all genomic markers reached the unadjusted p-value threshold of <5.00E-08.

However, given the over-conservative nature of the Bonferroni correction and also assuming that polygenic traits are likely to be dominated by numerous alleles with small causal effect we also report suggestive associations reaching the unadjusted 5.00E-08 p-value threshold (Table 1). Generally, two linear distances (nasal width and nasal tip protrusion), two angular distances (nasolabial angle and transverse nasal prominence angle), three indices (cephalic index, nasal index and nose-face width index) and one principal component revealed significant associations with 34 SNPs in 28 genes and intergenic regions (Table 1).

These factors can be arbitrarily divided into three main categories based on their cellular function: 1) genes with known roles in the craniofacial morphogenesis and/or mutated in various hereditary syndromes displaying craniofacial abnormalities; 2) genes or pseudo-genes without known function in the craniofacial morphology regulation or previously uncharacterized genes; and 3) non-protein coding genes, such as lncRNA class genes. There are also a number of significant variants that are located in the intergenic regions, with or without proximity to open reading frames (ORFs).

The majority of associated markers (n=21) are located in 17 protein-coding genes and pseudo-genes such as *AGXT2*, *CACNB4*, *COL11A1*, *EGFR*, *EYA1*, *EYA2*, *FAM49A*, *FOXN3*, *LHX8*, *LMNA*, *MYO5A*, *PAX3*, *PCDH15*, *RTTN*, *SMAD1*, *XXYLT1 and ZEB1*.

Five variants are present in RNA-coding (lncRNA) genes, which include *AC073218.1*, *RP11-494M8.4*, *RP11-408B11.2*, *RP11-785H20.1 and TEX41*.

The rest of the markers (n=6) are found in the intergenic regions, near the following genes and pseudogenes: *BMP4*, *HAS2-AS1*, *LOC124685*, *LOC100131241*, *MRPS36P3*, *PTCH1*, *SLC25A5P2*, *SV2B and TRNAY16P*.

Analysis of the functional annotation of significant markers revealed that one SNP represent missense mutation (rs37369), one SNP is a synonymous transversion (rs2290332), 21 markers are located in intronic sequences and 11 markers are located in intergenic regions (Table 1). The majority of significantly associated SNPs (n=27) are found in the regulatory elements of the genome, such as in transcription factor (TF) binding sites, and represent potentially functional SNPs (pfSNPs). These variants may be involved in “fine tuning” of the normal craniofacial phenotype as part of the enhancer/silencer mechanisms, as has been recently suggested [77].

The nasal area measurements, using either “n”, “prn”, “sn” or “al” landmarks, produced the majority of the total number of significant associations (6 out of 8). These measurements include nasal width (al-al), nasal tip protrusion (sn-prn), nasolabial angle (prn-sn-ls), transverse nasal prominence angle (t_l-prn-t_r), nasal index (al-al/n-sn), and nose face width index (al-al/zy-zy). The apparent overrepresentation of associations with the nasal area may be a result of the easier allocation and consequent superior reproducibility of the nasal area landmark measurements on 3D images. It may also be the result of specific selection of candidate genes from the JAX mice database resource, which focused on mutants that displayed various nasal area abnormalities.

The analysis of direct cranial measurements and their relative indices revealed significant associations only the Cephalic index (CI) with 9 SNPs.

The association analysis of the principal components (PC) representing all the craniofacial measurements, revealed one principal component (explaining 73.3% of all the craniofacial phenotypes) that was associated with 14 genetic markers (Table 1).

In contrast to most other craniofacial association studies that focused on a specific homogeneous population group (mostly Europeans), this study included samples from several population groups, which enabled investigation of the genetic factors influencing normal craniofacial morphology in different ethnicities [71]. Self-reported ancestry however, cannot be considered fully reliable, as demonstrated previously [72, 73]. In order to address this issue we assessed the self-reported ancestry using STRUCTURE with 186 SNPs removed due to long-range disequilibrium [49]. Following the rationale that the best ancestry estimates are obtained using a large number of random markers [74], we used all the available markers (after MAF filtering) in STRUCTURE analysis. The STRUCTURE analysis resulted in clusters of 367 Europeans, 51 East Asians, 43 South Asians and 16 Africans, with 107 samples designated as admixed ancestry (Fig. 1). Of the samples tested with STRUCTURE, 459 (89%) were assigned the same ancestry cluster (sole or mixed origin) as the self-reported information. Of the remaining 57 individuals, 39 were estimated as ‘admixture’ (based on up to 20% admixture threshold) and 18 were assigned a single ancestry, different to the self-reported ancestry (Fig. 1).

**Figure 1.**
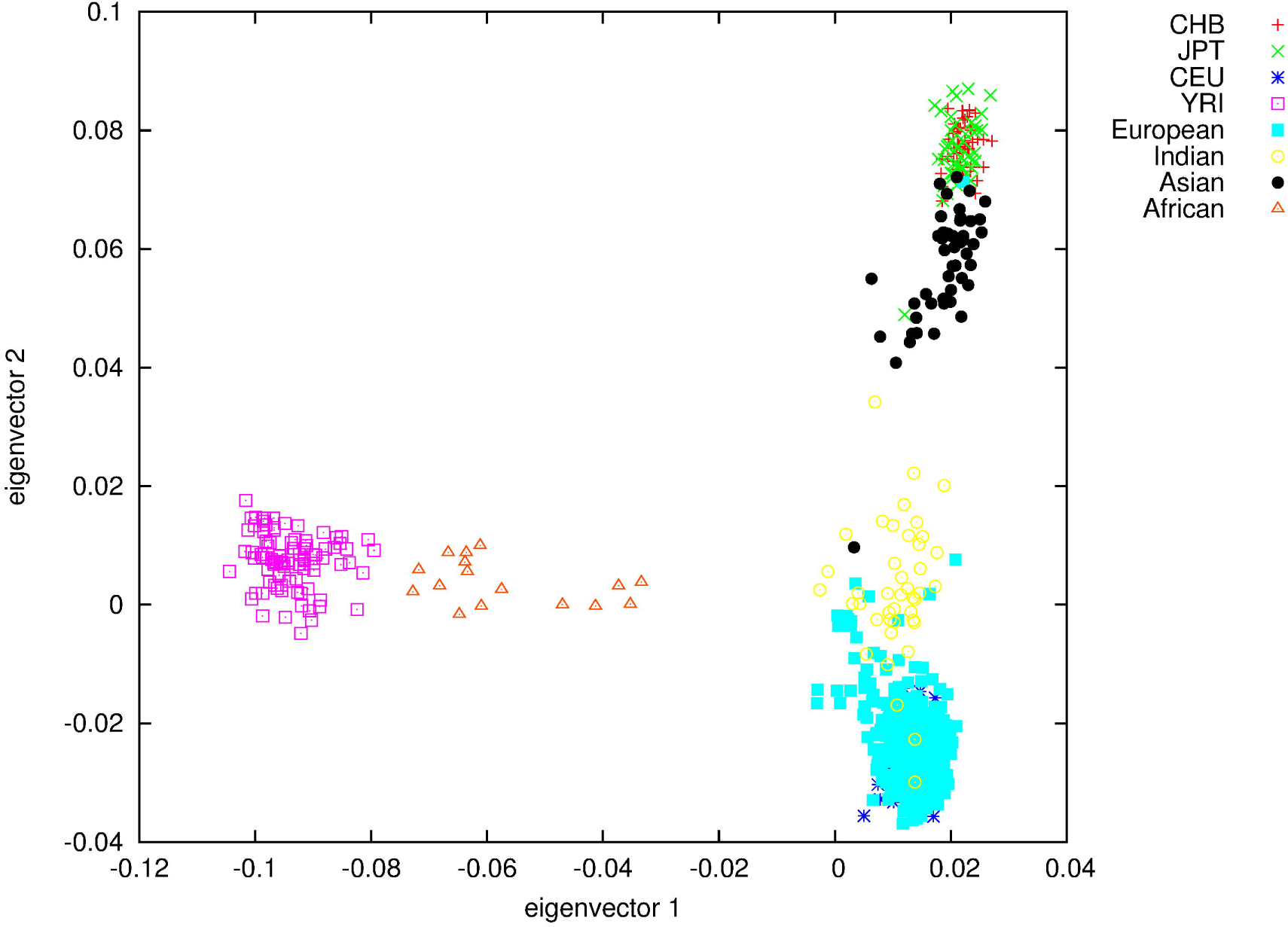
Anatomical position of the 32 manually annotated anthropometric landmarks used for calculation of linear and angular distances and ratios between the linear distances. Some landmarks are not clearly visible due to image orientation. gn = Gnathion, pg= Pogonion, sl = Sublabiale, li = Labiale Inferius, sto = Stomion, ls = Labiale superius, ch-r = Chelion right, ch-l = Chelion left, go-r = Gonion Right, go-l = Gonion left, sn = Subnasale, prn= Pronasale, al-r = Alare right; al-l= Alare left, n = Nasion, g= Glabella; tr = Tragion, en-l = left Endocanthion, en-r = right Endocanthion, ex-r = Right Endocanthion; ex-l = left Endocanthion, ps-r = Palpebrale superius right, ps-l = Palpebrale superius left, pi-r = Palpebrale inferius right, pi-l = Palpebrale inferius left, zy-r = Zygion Right, zy-l = Zygion Left, pra-r = Tragion right, pra-l = Tragion Left, sba-l = Subalare left, sa-l = Superaurale Left, pa-l = Postaurale left.

The risk of detecting false positive results because of population stratification was carefully assessed and further reduced by applying an EIGENSTRAT correction. Specifically, EIGENSTRAT’s smartpca.perl was used to perform PCA-clustering in comparison to reference populations from HapMap reference clusters. The Q-Q plots of the associated traits showed the expected distribution of data after applied correction (Supplemental Figs. S1-S8).

We did not perform allele imputations on this dataset because it includes individuals from heterogenous ancestral backgrounds, with 107 subjects classified as ‘admixture’, based on the applied threshold of 20%. Imputation using homogenous reference populations would have introduced unnecessary bias with wrongly imputed alleles in subsequent analysis steps.

#### Association analyses of the non-craniofacial traits

In our attempt to identify genetic markers influencing normal variation in craniofacial traits, we incorporated 522 markers previously associated with human pigmentation traits, such as eye, skin and hair colour. These markers were included to validate the statistical methods used for the craniofacial traits association study. The association analyses of the pigmentation traits, which were based on the HWE non-filtered data, did indeed confirm previously published findings, as detailed in Table S2. It should be noted however, that these results may not necessarily confirm the validity of the craniofacial markers associations.

The application of the Hardy-Weinberg equilibrium (HWE) threshold resulted in filtering 25% of the total number of SNPs. These markers included almost all the SNPs, previously associated with pigmentation traits, such as rs12913832, rs1129038, rs8039195 and rs16891982. This is not surprising, since population-related markers are likely not being in HWE ‘a priori’. Another explanation for this observation is potential bias from partially uncorrected heterogeneous ancestry, since the ancestry correction algorithm can only minimize, rather than completely remove spurious associations [52]. In fact, the association analyses of the HWE non-filtered genotyping data with pigmentation traits (eye, skin and hair colour), demonstrated highly significant associations, concordant with the literature (Table S2).

### Craniofacial gene and SNP annotations

The following section summarizes the genetic association results, providing brief annotation of the significantly associated genes and SNPs. Functional annotations, such as predicted molecular function, link to a biological process and a protein class of the 23 protein-coding genes and pseudo-genes (*AGXT2*, *BMP4*, *CACNB4*, *COL11A1*, *EGFR*, *EYA1*, *EYA2*, *FAM49A*, *FOXN3*, *LHX8*, *LMNA*, *MYO5A*, *PAX3*, *PCDH15*, *RTTN*, *SMAD1*, *XXYLT1 and ZEB1)* have been visualised using the PANTHER resource [78] and summarized in supplemental materials (Supplemental Figs. S17-S19).

#### Significantly associated genes with previously demonstrated role in craniofacial morphogenesis and/or mutated in hereditary syndromes displaying craniofacial abnormalities

A potentially functional SNP rs2289266 in the intron of the Paired Box 3 gene (*PAX3*) was associated with the transverse nasal prominence angle (p-value 1.95E-08). This gene is a member of the paired box (*PAX*) family of transcription factors, which play critical roles during foetal development. The *PAX3* protein regulates cell proliferation, migration and apoptosis. Mutations in *PAX3* are associated with Waardenburg syndrome (OMIM: 193500), which is characterized by a prominent and broad nasal root, a round or square nose tip, hypoplastic alae, increased lower facial height and other craniofacial abnormalities.

Notably, three other SNPs in this gene, rs974448, rs7559271 and rs1978860, were previously associated with normal variability of the nasion position [22] and the distance between the eyeballs and the nasion [20]. None of these SNPs were included in this study, as a result of primer design failure. No LD between rs2289266 and any of the previously associated markers in the *PAX3* gene was detected. Nevertheless, the association of another variant in the *PAX3* gene can be considered an independent confirmation of this gene’s involvement in regulation of normal craniofacial morphology.

SNPs rs4908280 and rs11164649 which are located in the regulatory element of the Collagen gene (*COL11A1*) intronic sequence, were associated with the cephalic index (p-values 1.66E-09 and 1.70E-08 respectively). *COL11A1* encodes one of the two alpha chains of type XI fibrillar collagen and is known to have multiple transcripts as a result of alternative splicing. The secreted protein is hypothesised to play an important role in fibrillogenesis by controlling lateral growth of collagen II fibrils.

Notably, the same variant (rs11164649) was recently linked to normal-range effects in various craniofacial traits, specifically eyes, orbits, nose tip, lips, philtrum and lateral parts of the mandible, although the measurements of the cephalic index were not performed in this study [93]. Our findings should be considered as independent confirmation of *COL11A2* gene and its specific polymorphism rs11164649 involvement in shaping the normal craniofacial morphology.

According to the MGI database, transgenic mice with shortened *COL11A2* mRNA (the second alpha chain of type XI fibrillar collagen) display abnormal facial phenotypes, including a triangular face and shorter and dimpled nasal bones [30]. Interestingly, *COL11A1* and other Collagen family genes were found to be mutated in Stickler (OMIM: 604841) and Marshall Syndromes (OMIM: 154780). These two inherited disorders display very similar phenotypes and each is characterized by a distinctive facial appearance, with flat midface, very small jaw, cleft lip/palate, large eyes, short upturned nose, eye abnormalities, round face and short stature. However, the facial features of Stickler syndrome are less severe and include a flat face with depressed nasal bridge and cheekbones, caused by underdeveloped bones in the middle of the face. Another member of the collagen family, *COL17A1*, was recently associated with the distance between the eyeballs and the nasion [23]. Our finding of genetic associations of additional members of the Collagen family provides further evidence of the importance of polymorphisms in these genes in determining the normal variety of specific craniofacial features.

Intergenic SNP rs942316, which is located upstream to the Bone Morphogenetic Protein (upstream to *BMP4*) gene, was strongly associated with the PC1phenotype (p-value 2.66E-09). The *BMP4* gene is a transforming growth factor, belonging to the beta superfamily, which includes large families of growth and differentiation factors. This gene plays an important role in the onset of endochondral bone formation in humans, including induction of cartilage and bone formation and specifically tooth development and limb formation. Gene onthology annotations related to this gene include heparin binding and cytokine activity. *BMP4* mutations have been associated with a variety of bone diseases, including orofacial cleft 11 (OMIM: 600625), Fibrodysplasia Ossificans (OMIM: 135100) and microphthalmia syndromic 6 (OMIM: 607932).

SNP rs2290332 represents a synonymous variant in the *Myosin VA* (Heavy Chain 12, *Myoxin*) gene (*MYO5A*). This variant was associated with the cephalic index (p-value 5.56E-10).

*MYO5A* is one of three myosin V heavy-chain genes, belonging to the myosin gene superfamily. *Myosin* V is a class of actin-based motor proteins involved in cytoplasmic vesicle transport and anchorage, spindle-pole alignment and mRNA translocation. It mediates the transport of vesicles to the plasma membrane, including melanosome transport. Mutations in this gene were associated with a number of neuroectodermal diseases, such as Griscelli syndrome. Additional mutations in this gene were associated with a rare inherited condition Piebaldism (OMIM:172800). The symptoms of Piebaldism include partial albinism and anomalies of the mouth area development, such as lips and philtrum abnormalities. Despite being a “silent” mutation, rs2290332 is located in the POLR2A TF binding site and may therefore affect various processes such as transcription, translation, splicing and mRNA transport, as has been shown in other studies [79].

Variant rs12041465, which is located in the intron of *LIM Homeobox 8 (LHX8*) was associated with transverse nasal prominence angle (p-value 2.30E-08). *LHX8* is a transcription factor and a member of the *LIM homeobox* family of proteins, which are involved in patterning and differentiation of various tissue types. Mutations in this gene were associated with clefts of the secondary palate in mouse model [80, 81].

Three intronic SNPs in the Eyes Absent Homolog 1 (*EYA1*) gene were associated with several craniofacial traits. The variant rs79867447 was associated with the nose width (p-value 3.92E-08), nasal index (p-value 3.53E-08), nose-face width index (p-value 1.10E-09) and PC1 (p-value 7.46E-10). The variant rs1481800 was associated with the cephalic index (p-value 2.07E-09). The variant rs73684719 was found in association with PC1 (p-value 2.62E-08). All three variants belong to potentially regulatory elements of the genome and are likely to affect TF binding sites. No linkage disequilibrium has been detected between these markers.

The *EYA1* encoded protein functions as histone phosphatase, regulating transcription during organogenesis in kidney and various craniofacial features such as branchial arches, eye and ear. *eya1* mutated mice display various craniofacial anomalies of the inner ear, mandible, maxilla and reduced skull [30]. Mutations in the human ortholog have been associated with several craniofacial conditions such as otofaciocervical syndrome (OMIM:166780), Weyers acrofacial dysostosis (OMIM:193530) and branchiootic syndrome (OMIM:608389).

Intronic SNP rs58733120 was associated with the nose width (p-value 5.37E-10), nasal index (p-value 9.46E-09), nose-face width index (p-value 3.38E-09) and PC1 (p-value 2.19E-08) phenotypes. This variant is located in the regulatory element of the *EYA2* gene, which belongs to the same eyes absent protein family as *EYA1* and plays a similar role in the embryonic development. An orthologue *eya2* gene encodes a transcriptional activator in mice and may play a role in eye development. Both *EYA1* and *EYA2* genes were shown to be expressed in the ninth week of human embryonic development [82]. None of the human craniofacial disorders were associated with *EYA2* gene to date.

SNP rs12076700 in the intron of the *Lamin A* gene (*LMNA*) was associated with the transverse nasal prominence angle (1.54E-09) and PC1 (p-value 7.87E-10).

*LMNA*, together with other *Lamin* proteins, is a component of a fibrous layer on the nucleoplasmic side of the inner nuclear membrane, which provides a framework for the nuclear envelope and also interacts with chromatin. *LMNA* encoded protein acts to disrupt mitosis and induces DNA damage in vascular smooth muscle cells, leading to mitotic failure, genomic instability, and premature senescence of the cell. This gene has been found mutated in Mandibuloacral Dysplasia which is characterized by various skeletal and craniofacial abnormalities, including delayed closure of the cranial sutures and undersized jaw [83].

Variant rs74884233 was associated with the transverse nasal prominence angle (p-value 1.20E-08). This variant is located in the intron of the *Rotatin* gene (*RTTN*). *RTTN* gene is involved in the maintenance of normal ciliary structure, which in turn effects the developmental process of left-right organ specification, axial rotation, and perhaps notochord development.

SNP rs17020235 was associated with the nasolabial angle (p-value 2.07E-09). This potentially functional variant is located in the intron of the *SMAD* Family Member 1 gene (*SMAD1*). *SMAD1* is a transcriptional modulator activated by BMP (bone morphogenetic proteins) type 1 receptor kinase, which is involved in a range of biological activities including cell growth, apoptosis, morphogenesis, development and immune responses.

*SMAD1* mutant mice display anterior truncation of the head with only one brachial arch present. In human, *SMAD1* mutations (together with *RUNX2*), are associated with the Cleidocranial Dysplasia (OMIM:119600), which is a Craniosynostosis-type disorder affecting cranial bones, palate and other tissues.

SNP rs950257 was associated with the PC1 trait (p-value 3.94E-08). This intronic variant is located in the *XXYLT1* gene, which codes for Xyloside Xylosyltransferase 1. This protein is an Alpha-1,3-xylosyltransferase, which elongates the O-linked xylose-glucose disaccharide attached to EGF-like repeats in the extracellular domain of Notch proteins signalling network. Notch proteins are the key regulators of embryonic development, which demonstrate a highly conserved sequence in various species. Interestingly, mutations in Notch proteins are associated with Hajdu–Cheney syndrome (OMIM:10250) and Alagille syndrome (OMIM:118450). The main phenotypic symptoms of these conditions include various malformations of the craniofacial tissues, including broad, prominent forehead, deep-set eyes and a small pointed chin.

SNP rs17335905 was associated with the nose-face width index (p-value 4.74E-08). This potentially functional variant is located in the intron of the *EGFR* gene, which encodes the Epidermal Growth Factor Receptor. *EGFR* is a cell surface protein that binds to epidermal growth factor (*EGF*). Binding of the protein to a ligand induces activation of several signalling cascades and leads to cell proliferation, cytoskeletal rearrangement and anti-apoptosis. Mouse carrying mutations in *EGFR*, express short mandible and cleft palate.

#### Significantly associated SNPs, located in genes or pseudo-genes that were not linked to craniofacial morphology regulation or genes with unknown function

Intronic variant rs59037879 in the Zinc Finger E-Box Binding Homeobox 1 (*ZEB1*) was found associated with cephalic index (p-value 6.27E-10), and transverse nasal prominence angle (p-value 5.31E-12). This gene encodes a zinc finger transcription factor, which is a transcriptional repressor. It regulates expression of different genes, such as interleukin-2 (*IL-2*) gene, ATPase transporting polypeptide (*ATP1A1*) gene and E-cadherin (*CDH1*) promoter in various cell types and also represses stemness-inhibiting microRNA. Mutations in this gene were previously associated with Corneal Dystrophy and various types of cancer.

A missense mutation rs37369 in the Alanine--Glyoxylate Aminotransferase 2 gene (*AGXT2*) was associated with nose width (p-value 1.04E-09), nasal index: (p-value 1.25E-10), transverse nasal prominence angle (p-value 1.46E-09) and PC1 (p-value 2.49E-11). This protein plays an important role in regulating blood pressure in the kidney through metabolizing asymmetric dimethylarginine (*ADMA*), which is an inhibitor of nitric-oxide (NO) synthase.

An intronic SNPs rs16830498, located in the regulatory element of the Calcium Channel Voltage-Dependent Beta 4 Subunit (*CACNB4*) gene intron, were significantly associated with cephalic index (p-value 7.57E-11).

The beta subunit of voltage-dependent calcium channels may increase peak calcium current by shifting the voltage dependencies of activation and inactivation, modulating G protein inhibition and controlling the alpha-1 subunit membrane targeting. *CACNB4* may be expressed in different isoforms through alternative splicing. Certain mutations in this gene have been associated with various forms of epilepsy, although no association with normal or abnormal craniofacial variation has been previously reported.

Potentially functional intronic SNP rs10825273 located in the regulatory elements of the Protocadherin-Related 15 (*PCDH15*) gene, was found in association with cephalic index (p-value 9.93E-09) and PC1 (p-value 1.04E-09). *PCDH15* is a member of the cadherin superfamily, which encodes an integral membrane protein that mediates calcium-dependent cell-cell adhesion and is known to have numerous alternative splicing variants. It plays an essential role in the maintenance of normal retinal and cochlear function. Mutations in this gene result in hearing loss and are associated with Usher Syndrome Type IIA (OMIM:276901).

Two intronic variants in the Family With Sequence Similarity 49 Member A gene (*FAM49A*) were associated with multiple craniofacial traits. rs6741412 was found in association with the transverse nasal prominence angle (p-value 2.75E-09) and PC1 (p-value 4.67E-10). rs11096686 was associated with PC1 (p-value 2.25E-08). The *FAM49A* protein is known to interact with hundreds of miRNA molecules during pre-implantation of the mouse embryo and also expressed in the developing chick wing, but no information on its specific function or disease association have been identified.

SNP rs390345, located in the intronic regulatory sequence of the Forkhead Box N3 gene (*FOXN3*), was associated with the PC1 (p-value 7.46E-11). *FOXN3* encodes multiple splicing variants and acts as a transcriptional repressor. It is proposed to be involved in DNA damage-inducible cell cycle arrests at G1 and G2. There are no previous reports on *FOXN3* association with either normal craniofacial development or pathological conditions.

#### Significantly associated SNPs located in the non-protein coding genes, such as lncRNA class genes

Intronic SNP rs10496971 in the *TEX41* (Testis Expressed 41) gene produced significant associations with transverse nasal prominence angle (p-value 5.52E-09), cephalic index (p-value 5.315E-09) and PC1 (p-value 3.71E-09).

*TEX41* is a long intergenic non-protein coding RNA (lncRNA) class gene, which is located on chromosome 2 and has 43 transcript variants as a result of alternative splicing. lncRNAs are known as regulators of diverse cellular processes. However, the function of this gene remains unknown. Despite its name, this gene is expressed in a variety of tissues, with the highest demonstrated levels in kidney. Its potential involvement in craniofacial genetics, and specifically in influencing normal facial variation, has not been reported previously. Notably, the rs10496971 variant is located in the regulatory element of the genome (as well as 49 other associated SNPs) and may influence normal craniofacial morphology by affecting either enhancer or silencer sequences or transcriptional factor (TF) binding sites [77].

The SNP rs1482795, located in the RNA gene *RP11-494M8.4*, was associated with the nose width (p-value 7.68E-10) and nasal index (p-value 1.83E-08) measurements.

Both SNPs rs892457 and rs892458 located in the non-protein coding lncRNA gene *AC073218.1*, were associated with the transverse nasal prominence angle (p-value 3.43E-08) and (p-value 1.73E-08), respectively.

SNP rs7311798, located in the lncRNA gene *RP11-408B11.2* was associated with the nasal index (p-value 1.77E-08).

SNP rs7844723 in the *RP11-785H20.1* (lncRNA gene) was associated with the PC1 (p-value 8.11E-09) phenotype.

SNP rs2357442 was associated with the transverse nasal prominence angle (p-value 4.40E- 08). This variant is located in the Long Interspersed Nuclear Element 1 (*LINE-1*) retrotransposon sequence, which in turn shows homology with uncategorized mRNA KC832805 on the Y-chromosome.

*LINE-1* elements comprise approximately 21% of the human genome, and have been shown to modulate expression and produce novel splice isoforms of transcripts from genes that span or neighbour the LINE-1 insertion site. In addition, rs2357442 is located close to three pseudo-genes with unknown function: *SLC25A5P2*, *LOC100130842* and *RP11-1033H12.1*, while the last two represent RNA-coding lncRNA genes.

#### Significantly associated SNPs located in the intergenic regions

SNP rs10512572, located between *Serpine1 MRNA Binding Protein 1* pseudogene (*LOC100131241*) and *MyosinLight Chain 6 Alkali Smooth Muscle and Non-Muscle* pseudogene (*LOC124685*), was associated with nasal tip protrusion (p-value 2.22E-08), transverse nasal prominence angle (p-value 1.38E-11) and PC1 (p-value 4.99E-10). While pseudogenes in general are non-protein coding, their sequences can be functional and play important roles in different biological processes [85]. It should be noted that some genes may be incorrectly defined as pseudogenes, based solely on their sequence computational analysis [86]. The function of these two pseudogene sequences is unknown.

SNP rs8035124 was significantly associated with the nose width (p-value 1.52E-10). This variant is located between the Synaptic Vesicle Glycoprotein 2B (*SV2B*) and Transfer RNA Tyrosine 16 (Anticodon GUA) Pseudogene (*TRNAY16P*) genes. The *SV2B* is a protein coding gene, which plays a role in the control of regulated secretion in neural and endocrine cells. The *TRNAY16P* is a pseudogene with unknown function.

Additional SNP rs373272 was associated with cephalic index (p-value 2.40E-08). However, no genes were identified within 50 kb window of its chromosomal location.

## Discussion

This study focused on the identification of genetic markers in a set of candidate genes associated with various craniofacial traits, representing the most comprehensive scan for genetic markers involved in normal craniofacial development performed to date. We identified 8 craniofacial significantly associated (unadjusted p-value < 5.00E-08) with 34 genomic variants in 28 genes and intergenic regions. Following the application of Bonferroni correction (adjusted p-value threshold of 1.6E-07), associations were observed between 5 craniofacial traits (nasal width, cephalic index, nasal index, transverse nasal prominence angle and principal component) and 6 SNPs (rs8035124, rs16830498, rs37369, rs59037879, rs10512572 and rs390345) located in 6 genes and intergenic regions (15q26.1, 17q24.3, *CACNB4*, *AGXT2*, *ZEB1 and FOXN3* respectively). We report all the significant markers that met the less stringent GWAS threshold (p-value<5.00E-08), as Bonferroni correction is generally considered over-conservative, especially when analysing complex traits such as craniofacial morphology, which is likely to be influenced by a large number of alleles with relatively small individual effect, similar to height [89, 90].

The association of the *PAX3* gene and the *COL11A1* gene with transverse nasal prominence angle and cephalic index respectively, confirms previous findings [11, 22, 23, 91]. In fact, an intronic SNP rs11164649 that was associated with cephalic index in the current study, was recently associated with normal-range effects in various craniofacial traits and used for their prediction [91], while the other variants in *COL11A1* (rs4908280) and in *PAX3* (rs2289266) have not been reported previously. The rest of the identified associations are also novel. These include 21 significantly associated markers in protein-coding genes and pseudo-genes, such as *AGXT2*, *CACNB4*, *EGFR*, *EYA1*, *EYA2*, *FAM49A*, *FOXN3*, *LHX8*, *LMNA*, *MYO5A*, *PCDH15*, *RTTN*, *SMAD1*, *TEX41*, *XXYLT1 and ZEB1*. Additional 7 significantly–associated SNPs are found in intergenic regions adjacent to several loci, such as *BMP4*, *LOC124685*, *LOC100131241 and PTCH1*. Some of these genes were previously linked to craniofacial embryogenesis, while others represent novel associations.

Six genetic variants were found in lncRNA genes, which have not been previously linked to craniofacial morphogenesis before. These findings may suggest there may be a yet unexplored level of epigenetic regulation affecting craniofacial morphology. lncRNAs are a recently discovered class of factors, whose expression is thought to be important for the regulation of gene expression through several different mechanisms involving competition with transcription by recruitment of specific epigenetic factors to promoter regions, as well as indirectly affecting gene expression by interacting with miRNA and other cellular factors [92]. The comprehensive role of epigenetic regulation in general, and in craniofacial embryonic development in particular, is poorly understood. There is a limited number of recent studies revealing thousands of enhancer sequences, predicted to be active in the developing craniofacial complex in mice [77, 93] and potentially in humans. Both the epistatic and epigenetic interactions may represent a more complex level of craniofacial morphology regulation and require further investigation.

Even though a relatively high number of phenotypes were studied (92 linear and angular measurements and indices), this may still represent an oversimplification of the complexity of the human face. Despite the importance of the association between specific 3D measurements and SNPs demonstrated in this study, the association of facial shapes, represented by the principal components should better represent the face. Given that embryonic developmental processes such as cell proliferation, polarity orientation and migration occur in a 3D environment, principal components that in essence denote specific facial shapes, may provide a more accurate representation of these processes. However, only one of the 10 principle components showed significant associations at the GWAS threshold level. While the explanation of this observation is unclear, it is consistent with other similar studies [22, 23]. The specific anthropometric measurements on the other hand, produced numerous significant associations, identifying many genes and intergenic regions that appear to play important roles in the development of normal human facial appearance. The major limitation of this study is the replication of these results that has not been performed yet due to time and budget constraints. However, the confirmation of the two previously associated genes (*PAX3* and *COL11A1*) supports the validity of our findings.

Given the high complexity of the face, as well as the composite nature of the genetic regulation that affects its development, alternative comprehensive approaches of capturing facial morphology would be beneficial. A number of such methods has recently revealed additional genes with specific polymorphisms associated with the development of craniofacial traits within the normal variation range [91, 94]. Further studies may involve the use of these or alternative methods to capture the majority of variation in craniofacial traits. Craniofacial phenotypes, together with additional external visible traits such as sex, age and BMI and ancestry, could be treated as a “vector”, which could then be used to predict appearance [95].

A recent attempt to predict facial appearance was performed using only 24 SNPs [96]. This approach has promise, although it is largely based on reconstruction of a ‘facial composite image’ through prediction of ancestry, sex, pigmentation and human perception of faces. This approach is reasonable, but it does not negate the use of association studies looking at specific craniofacial traits. Genetic association studies of a large scope of individual anthropometric measurements are essential to provide information on specific genes and their polymorphisms, which affect these traits and may therefore be useful in predicting the size and the shape of specific facial features.

Additional association studies on large sample sizes, incorporating dense SNP panels or whole genome sequencing approaches, in conjunction with either a comprehensive set of anthropometrical measurements or morphologically adequate representation of the craniofacial characteristics would be a valuable adjunct to the promising results obtained in this study. These studies will not only improve our understanding of the genetic factors regulating craniofacial morphology, but will also enable a better prediction of the visual appearance of a person from DNA.

## Methods

### Sample collection and ethics statement

A total of 623 unrelated individuals, mostly Bond University (Gold Coast, Australia) students, of Australian ancestry were recruited. The participants provided their written informed consent to participate in this study, which was approved by the Bond University Ethics committee (RO-510). To minimize any age-related influences on facial morphology the samples were largely collected from volunteers aged between 18 and 40. The mean age of the volunteers was 26.6 (SD ± 8.9). Following the exclusion of the individuals who had experienced severe facial injury and/or undergone facial surgery (e.g. nose or chin plastics) 587 samples remained for the further step of DNA sequencing.

Each participant donated four buccal swabs (Isohelix, Cell Projects, Kent, UK). 3-Dimentional (3-D) facial scans and three direct cranial measurements were obtained as described below. Samples with low DNA quantity or low quality facial scans were eliminated leaving 587 DNA samples for subsequent genotyping.

Additional phenotypic trait information such as height, weight, age, sex, self-reported ancestry (based on the grandparents from both sides), eye lid (single or double), ear lobe (attached or detached), hair texture (straight, wavy, curly or very curly), freckling (none, light, medium or extensive), moles (none, few or many), as well as eye skin, and hair pigmentation was collected by a single examiner in order to reduce potential variation. The pigmentation traits were arbitrary assigned according to previously published colour charts [26–28].

### 3D images collection and analysis

Craniofacial scans were obtained using the Vivid 910 3-D digitiser (Konica Minolta, Australia) equipped with a medium range lens with a focal length of 14.5 mm. The scanner output images were of 640 x 480 pixels resolution for 3D and RGB data. Two daylight fluorescent sources (3400K/5400K colour temperature) were mounted at approximately 1.2 meters from the subject’s head to produce ambient light conditions.

The scanner was mounted approximately one meter from the volunteer’s head. Each volunteer remained in an upright seated position and kept a neutral facial expression during the scan. Subjects with long hair pulled their hair behind the ears or were asked to wear a hair net. Glasses and earrings were removed.

Each volunteer was scanned from a distance of approximately one meter from three different angles (front and two sides). The final merged 3D image was produced by semi-automatically aligning the three scans and manually cropping non-overlapping or superfluous data such as the neck area and hair using Polygone^®^ software (Qubic, Australia). The complete coordinates of each merged 3D image were then saved in a ‘vivid’ file format (vvd) and exported to Geomagic^®^ software (Qubic, Australia) for subsequent image processing.

Based on the anthropometrical literature [29] 32 anthropometrical landmarks were manually identified on each 3-D image using the Geomagic software (Fig. 2 and Supplemental Table S1). Each landmark was represented by ‘x’, ‘y’ and ‘z’ coordinates as part of the Cartesian coordinate system. The coordinates were exported to an Excel spreadsheet for subsequent calculation of 86 Euclidean distances, including 54 linear distances, 10 angular distances and 21 indices (ratios) between the linear distances (Fig. 2 and Table 2).

**Figure 2.**
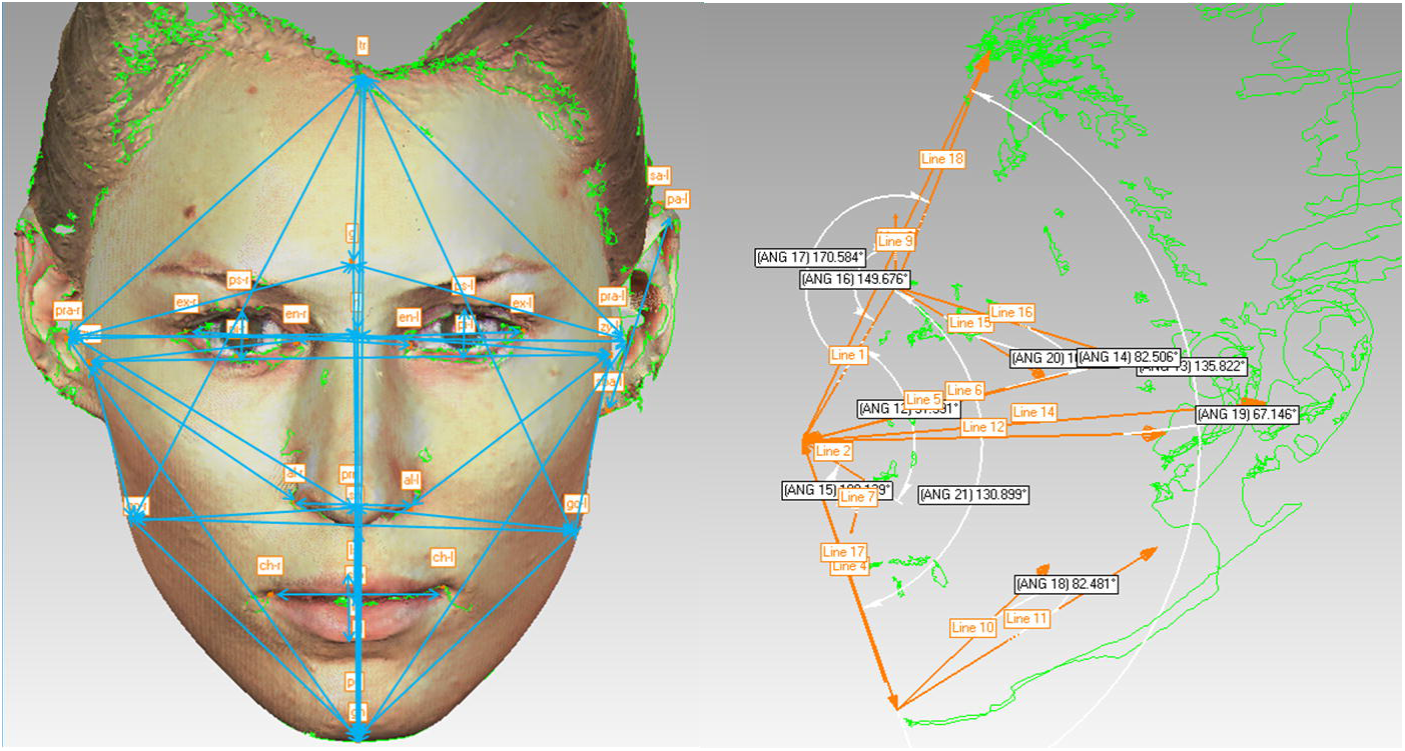
Illustration of linear and angular distances calculated from manually annotated landmark coordinates.

**Table 2.**
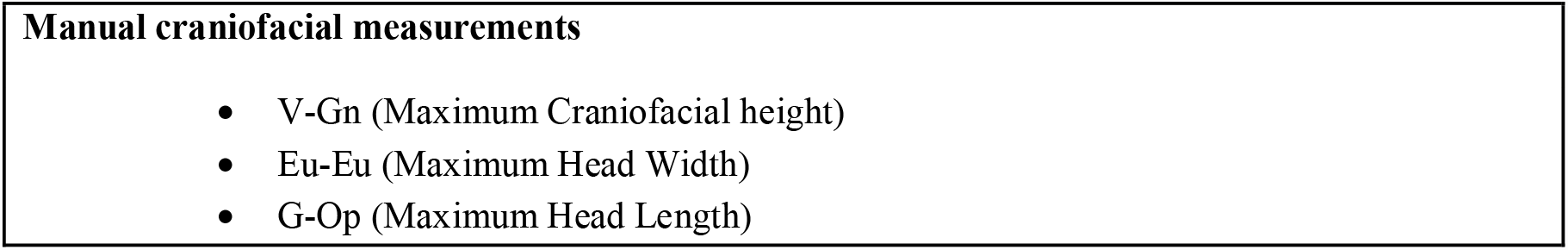
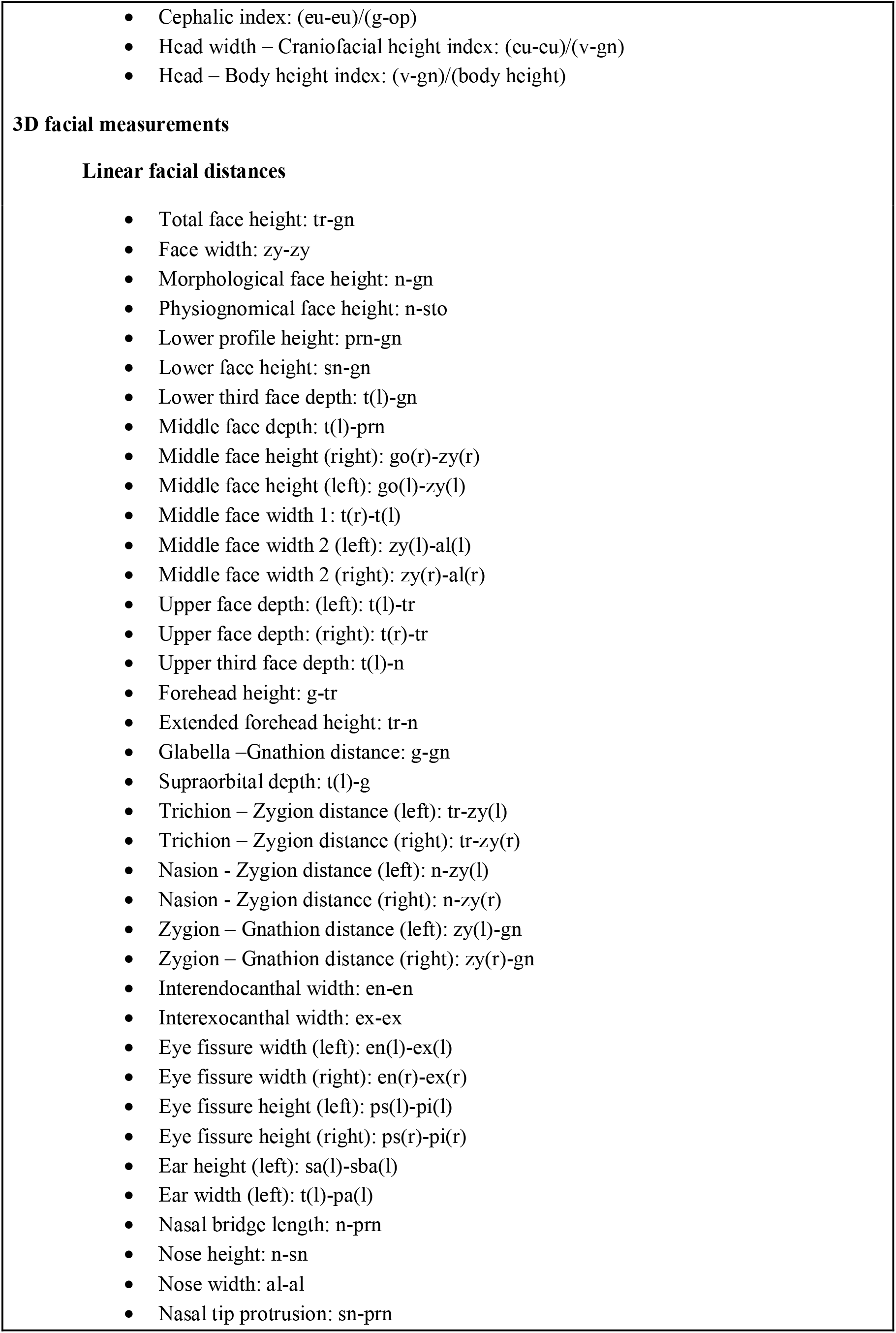
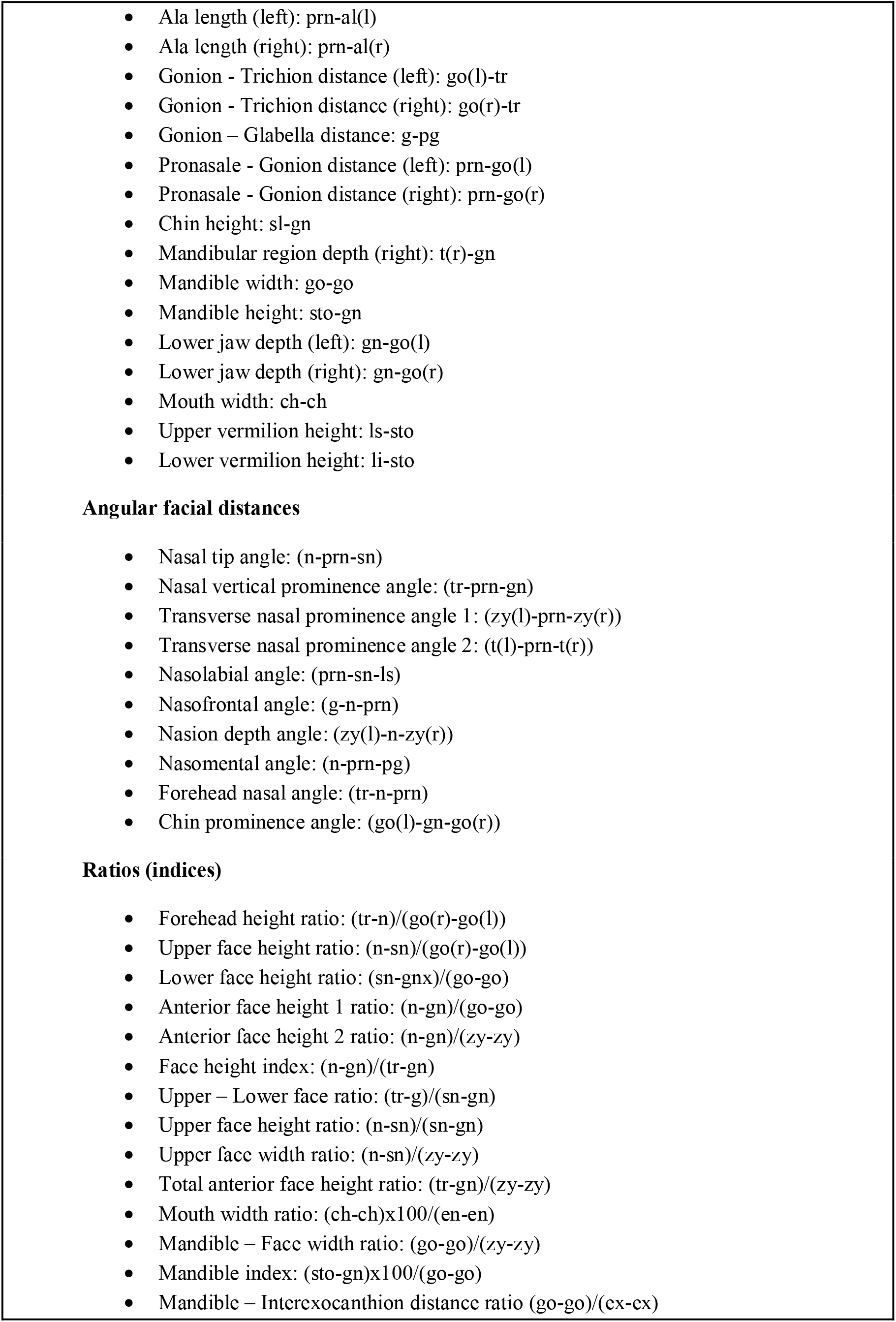
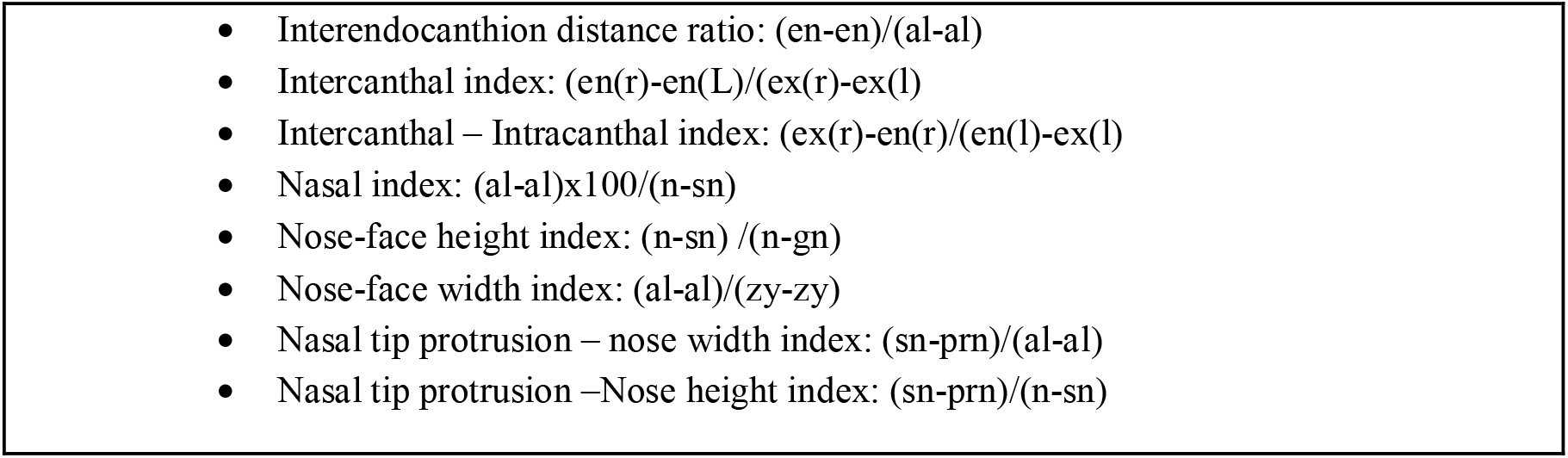
Craniofacial anthropometric measurements recorded in the study and used for genetic association analyses.

Additionally, three direct cranial measurements: maximum cranial breadth (Euryon – Euryon), maximum cranial length (Gonion – Opisthocranium) and maximum cranial height (Vertex – Gnathion), were collected manually using a digital spreading calliper (Paleo-Tech Concepts, USA). Based on the craniofacial and body height measurements, three craniofacial ratios were calculated: Cephalic index: (eu-eu)/(g-op), Head width – Craniofacial height index: (eu-eu)/(v-gn) and Head – Body height index: (v-gn)/(body height), as summarised in Table 2.

### Phenotypic traits summary

A total of 54 linear distances, 10 angular distances and 21 indices (ratios) between the linear distances were calculated based on the Cartesian coordinates of 32 anthropometric landmarks that were manually mapped on each of the 587 3-D facial images (Fig. 2, Fig. 3, Table 2 and Supplemental Table S1). Three additional craniofacial distances were obtained by direct measurement of subjects’ heads and used to calculate three indices: maximum cranial breadth, maximum cranial length and maximum cranial height, cephalic index, head width – craniofacial height index and head – body height index (Table 2). Information on the eyelid and earlobe morphology (single/double and attached/detached respectively) was recorded. Furthermore, the linear and angular facial distances were used to calculate 10 principal components (PCs). Additional phenotypic traits such as eye, skin and hair pigmentation, hair texture, freckling, moles, height, weight, BMI, age and sex were collected. In total, the data on 104 craniofacial phenotypic traits were recorded and used for genetic association analyses.

**Figure 3.**
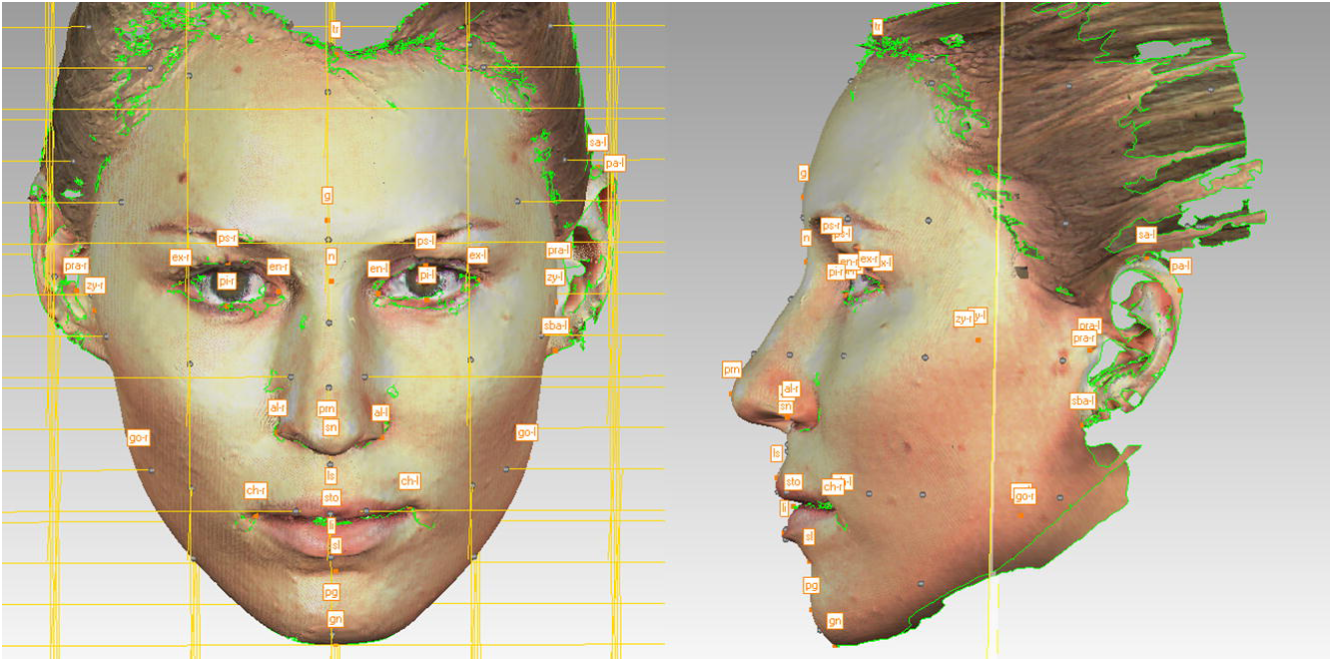
Population structure as represented by plotting genomic PCs 1 and 2, using 270 HapMap individuals as anchor clusters. YRI: Yoruba, Nigeria, Africa. JRI: Japanese, Tokyo, Japan. CHB: Han Chinese, Beijing, China. CEU: Utah residents with European ancestry.

The phenotypic data collection by a single examiner achieved more consistent measurements from the 3-D image analyses. In addition, all measurements were based on the images of participants within a narrow age range 26.6 (SD ± 8.9).

### DNA extraction and quantification

DNA was purified from buccal swabs using the Isohelix DDK isolation kit (Cell Projects, Kent, UK) according to the manufacturer instructions. DNA samples were quantified using a Real Time quantitative PCR (q-PCR) method using a Bio-Rad CFX96 (Bio-Rad, Gladesville, Australia). This assay amplified a 63bp region of the OCA locus. The primer sequences were 5′-GCTGCAGGAGTCAGAAGGTT-3′ (forward primer) and 5′-CATTTGGCGAGCAGAATCC-3′ (reverse primer) at a final concentration of 200mM. All DNA samples were additionally quantified using the Qubit 2.0 fluorimeter (Invitrogen) prior to library construction as per manufacturer recommendations.

### Candidate genes and SNPs selection

Two main complementary strategies were used to generate a preliminary list of candidate genes and genetic markers. The first focused on searching the literature and web resources for candidate genes involved either in normal craniofacial variation or in craniofacial malformations in humans and model organisms (Supplemental Table S2).

The search for candidate genes focused not only on specifically defined craniofacial disorders, but also on genetic syndromes with various manifestations of craniofacial malformations, such as Down syndrome, Noonan Syndrome, Floating-Harbor Syndrome and others, as detailed in Supplemental Table S2. The main resources for locating candidate genes in the animal models were Mouse Genome Informatics [30] and AmiGo tool [31] The main resources for identifying candidate genes in the human genome were OMIM [32] and GeneCards [33]. A comprehensive list of web resources used for candidate gene search is detailed in the Supplemental Appendix S1.

The second approach initially implemented a broad search for high Fst SNPs, such as ancestry informative markers (AIMs), with the rationale that many genes affecting craniofacial traits would have significantly different allele frequencies across populations. AIMs were selected from a variety of published and online resources [34–43].

The relevant genes obtained by both approaches were subsequently checked for potential involvement in craniofacial embryogenesis, limb development and bilateral body symmetry. It should be noted however, that the final candidate gene list was not limited to craniofacial genes and included high Fst SNPs in genes with unknown function as well as markers located in intergenic regions, potentially possessing regulatory functions.

The resulting set of SNPs was further screened for high Fst SNPs (≥0.45) in three ‘1000 genomes’ populations (CAU, ASW, CHB) using ENGINES browser [44] as well as potentially functional polymorphisms, such as non-synonymous SNPs [45], markers in transcription factor binding sites [46] and splicing sites [47] using various web resources, as detailed in Supplemental Appendix S1 and reviewed on the GenEpi website [48]. The candidate markers search resulted in identification of 1,319 SNPs, located in approximately 177 genes/intergenic regions, as discussed in the Results section.

The chromosomal locations of final candidate markers were submitted to the custom Ampliseq primer design pipeline (Life Technologies), according to manufacturer recommendations. There were primer design difficulties for 881 markers. The marker list was therefore redesigned to include alternative tagging markers showing high linkage disequilibrium with the markers that failed initial primer design, resulting in 1,670 candidate genetic markers. Inclusion of SNPs with MAF<1% added additional 4,381 genetic markers (6,051 in total). The final custom Ampliseq panel was manufactured as two separate pools of 849 and 847 primer pairs, with each amplicon covering between 125 bp and 225 bp, therefore possibly containing more than one polymorphism, and in total covering 15.78 kb of the reference human genome. This panel included 1,319 initially targeted craniofacial and pigmentation candidate markers as well as 4,732 markers in LD with original candidate SNPs that failed primer design.

Inclusion of novel, rare SNPs (MAF<1%) increased the final number of genotyped markers to 8,518 SNP in all sequenced DNA samples, although the markers with MAF≤2% were not included in the association study. The list of all genotyped markers and their respective genes is detailed in Table S1.

### SNP genotyping and data analysis

Multiple DNA libraries were constructed from sets of 32 Ion Xpress^TM^ (Life Technologies) barcoded samples using the Ion AmpliSeq^TM^ library Kit 2.0 (Life Technologies) in conjunction with two custom primer mixes that were pooled according to manufacturer recommendations. Libraries were quantified using the Ion Library Quantitation kit (Life Technologies) and pooled in equal amounts for emulsion PCR, which was performed using the OneTouch^TM^ 2 instrument (Life Technologies) according to manufacturer recommendations. 587 DNA samples were genotyped by massively parallel sequencing on the Personal Genome Machine (PGM) (Life Technologies) using the Sequencing 200 v2 kit and 316 Ion chips (Life Technologies).

Raw sequencing data were collected and processed on the Torrent Suite Server v3.6.2 using default settings. Alignment and variant calling were performed against the human genome reference (hg19) sequence at low stringency settings. Binary alignment map (BAM) files were generated and exported to the Ion Reporter^TM^ (IR) cloud-based software for SNP annotation against the reference hotspot file. The IR analysis resulted in generation of the individual variant caller files (VCF) with genotype calls for each sample as well as various statistics of the sequencing quality.

To reduce potential bias of the self-reported ancestry, ancestry inferences were obtained by 3,302 markers using STRUCTURE version 2.3.4 with default parameters as per software developer recommendations [49]. SNPs in long-range Linkage Disequilibrium (> 100,000 bp) were excluded from the STRUCTURE run. The ancestry was estimated based on four predefined population clusters: Europeans, East Asians, South Asians and Africans, according to software developer recommendations. Relative allele calls for four predefined HapMap population clusters (CEU, YRI, CHB and JPT) were used as reference populations [50]. The ancestry origin was estimated as a single (unmixed) source where the main ancestry cluster could be affiliated with at least 80% of the total mixed ancestry. The samples with mixed ancestry (>20% admixture) were assigned to an ‘Admixture’ cluster.

Association analyses were performed using SNP & Variation Suite v7 (SVS) (Golden Helix, Inc., Bozeman, MT) and replicated using PLINK v1.07 software [51]. Statistical analyses in both software programs were performed using linear regression with quantitative phenotypes, and logistic regressions with binary phenotypes under the assumption of an additive genetic model, while each genotype was numerically encoded as 0, 1 or 2. Population stratification correction, incorporated by EIGENSTRAT program was implemented in the analyses [52, 53]. In order to reduce any potential confounding effects, all the craniofacial traits association analyses were performed using sex, BMI and EIGENSTRAT ancestry clusters as covariates. In PLINK, p-values were adjusted using the ‘–adjust’ option. The final reported association results are based on the PLINK statistical analyses with the EIGENSTRAT PCA clusters, BMI and gender as covariates.

Annotation analysis of the significantly associated genes was performed using the GeneCards, ENTREZ and UniProtKB web portals [33, 54]. The MalaCards web site was used to detect association between the genes and hereditary syndromes [55]. The GeneMania web site was used to identify a functional network among the genes and encoded proteins [56]. Gene ontology web resource was used to find orthologs of human genes in other organisms [31, 57]. The MGI database was used to search for the phenotype in relevant craniofacial mouse gene mutants [30]. The dbSNP, 1000 genomes, SNPnexus and Alfred websites were used for SNP annotations [58–61].

The SNP Annotation and Proxy Search (SNAP) web portal was used to find SNPs in linkage disequilibrium (LD) and generate LD plots, based on the CEU population panel from the 1000 genomes data set, within a distance of up to 500kb and an r^2^ threshold of 0.8 [62].

The Regulome database and potentially functional database (PFS) searches were implemented to annotate SNPs with known and predicted regulatory elements in the intergenic regions of the *H. sapiens* genome [47, 63].

## List of abbreviations

3D: 3-Dimentional
AIMs: ancestry informative markers
ASW: African ancestry in Southwest USA
BAM: Binary alignment map
BMI: Body Mass Index
CAU: Caucasian
CHB: Han Chinese in Beijing, China
ENGINES: ENtire Genome INterface for Exploring Snps
EVT: Externally visible characteristic
DVI: Disaster victim identification
FDP: Forensic DNA phenotyping
GWAS: Genome wide association studies
HWE: Hardy-Weinberg equilibrium
JPT: Japanese in Tokyo, Japan
LD: linkage disequilibrium
lncRNAs: long non-coding RNAs
LINE-1: Long Interspersed Nuclear Element 1
MAF: Minor allele frequency; measurement error
ME: Measurement error
MD: Mean difference
OMIM: Online Mendelian Inheritance in Man
ORFs: open reading frames
pfSNP: Potentially functional SNP
PCA: Principal component analysis
RGB: Red, Green, Blue (colours)
SNP: Single-nucleotide polymorphism
SNAP: SNP Annotation and Proxy Search
STR: Short tandem repeat
TF: Transcription factor
VCF: Variant Call Format
YRI: Yoruba in Ibadan, Nigeria

## Declarations

### Ethics and consent to participate

The participants provided their written informed consent to participate in this study, which was approved by the Bond University Ethics committee (RO-510).

## Authors’ contributions

MB designed the study, carried out the molecular genetic studies, carried out the data analysis, participated in the statistical analysis and drafted the manuscript. PB performed the statistical analysis and drafted the manuscript. AvD participated in the design of the study and drafted the manuscript. All authors read and approved the final manuscript.

## Consent to Publish

Not applicable

## Availability of data and materials

The genomic data supporting the conclusions of this article are included within the article and its additional files.

## Acknowledgments

We would like to thank the volunteers who participated in this study without whom we could not have performed this research. We would like to thank Olga Kondrashova who helped with the sample collection. We also thank Technical Support Working Group (TSWG) and Pelerman Holdings Pt Ltd for their generous support of this project.

## Additional files

**Figure S1. Q-Q plot of the PCA-corrected -log10 p-values for the difference between the observed association for the tails of al-al distance and expected association based on the overall al-al distance distribution.**

**Figure S2. Q-Q plot of the PCA-corrected -log10 p-values for the difference between the observed association for the tails of sn-prn distance and expected association based on the overall sn-prn distance distribution.**

**Figure S3. Q-Q plot of the PCA-corrected -log10 p-values for the difference between the observed association for the tails of cephalic index and expected association based on the overall cephalic index distribution.**

**Figure S4. Q-Q plot of the PCA-corrected -log10 p-values for the difference between the observed association for the tails of nasal index and expected association based on the overall nasal index distribution.**

**Figure S5. Q-Q plot of the PCA-corrected -log10 p-values for the difference between the observed association for the tails of nose-face width index and expected association based on the overall nose-face width index distribution.**

**Figure S6. Q-Q plot of the PCA-corrected -log10 p-values for the difference between the observed association for the tails of nasolabial angle and expected association based on the overall nasolabial angle distance distribution.**

**Figure S7. Q-Q plot of the PCA-corrected -log10 p-values for the difference between the observed association for the tails of transverse nasal prominence angle and expected association based on the overall transverse nasal prominence angle distribution.**

**Figure S8. Q-Q plot of the PCA-corrected -log10 p-values for the difference between the observed association for the tails of PC1 trait and expected association based on the overall PC1 trait distance distribution.**

**Figure S9. Manhattan plot of the genomic associations of the al-al distance, based on the initial p-values from analysis of the PCA-corrected data. The -log10 (P value) is plotted against the physical positions of each SNP on each chromosome. The basic significance threshold is indicated by the blue line for -log10(1e-5) and the genome-wide significance threshold for - log10(5e-8) is indicated by the red line.**

**Figure S10. Manhattan plot of the genomic associations of the sn-prn distance, based on the initial p-values from analysis of the PCA-corrected data. The -log10 (P value) is plotted against the physical positions of each SNP on each chromosome. The basic significance threshold is indicated by the blue line for -log10(1e-5) and the genome-wide significance threshold for -log10(5e-8) is indicated by the red line.**

**Figure S11. Manhattan plot of the genomic associations of the cephalic index, based on the initial p-values from analysis of the PCA-corrected data. The -log10 (P value) is plotted against the physical positions of each SNP on each chromosome. The basic significance threshold is indicated by the blue line for -log10(1e-5) and the genome-wide significance threshold for -log10(5e-8) is indicated by the red line.**

**Figure S12. Manhattan plot of the genomic associations of the nasal index, based on the initial p-values from analysis of the PCA-corrected data. The -log10 (P value) is plotted against the physical positions of each SNP on each chromosome. The basic significance threshold is indicated by the blue line for -log10(1e-5) and the genome-wide significance threshold for - log10(5e-8) is indicated by the red line.**

**Figure S13. Manhattan plot of the genomic associations of the nose-face width index, based on the initial p-values from analysis of the PCA-corrected data. The -log10 (P value) is plotted against the physical positions of each SNP on each chromosome. The basic significance threshold is indicated by the blue line for -log10(1e-5) and the genome-wide significance threshold for -log10(5e-8) is indicated by the red line.**

**Figure S14. Manhattan plot of the genomic associations of the nasolabial angle, based on the initial p-values from analysis of the PCA-corrected data. The -log10 (P value) is plotted against the physical positions of each SNP on each chromosome. The basic significance threshold is indicated by the blue line for -log10(1e-5) and the genome-wide significance threshold for -log10(5e-8) is indicated by the red line.**

**Figure S15. Manhattan plot of the genomic associations of the transverse nasal prominence angle, based on the initial p-values from analysis of the PCA-corrected data. The -log10 (P value) is plotted against the physical positions of each SNP on each chromosome. The basic significance threshold is indicated by the blue line for -log10(1e-5) and the genome-wide significance threshold for -log10(5e-8) is indicated by the red line.**

**Figure S16. Manhattan plot of the genomic associations of the PC1 trait, based on the initial p-values from analysis of the PCA-corrected data. The -log10 (P value) is plotted against the physical positions of each SNP on each chromosome. The basic significance threshold is indicated by the blue line for -log10(1e-5) and the genome-wide significance threshold for -log10(5e-8) is indicated by the red line.**

**Figure S17. Pie chart, illustrating molecular function classification of human genes, harbouring genomic markers in significant association with craniofacial phenotypes. The genes include: *AGXT2*, *BMP4*, *CACNB4*, *COL11A1*, *EGFR*, *EYA1*, *EYA2*, *FAM49A*, *FOXN3*, *LMNA*, *MYO5A*, *PAX3*, *PCDH15*, *RTTN*, *SMAD1*, *XXYLT1 and ZEB1*.**

**Figure S18. Pie chart, illustrating biological processes classification involving human genes, harbouring genomic markers in significant association with craniofacial phenotypes. The genes include: *AGXT2*, *BMP4*, *CACNB4*, *COL11A1*, *EGFR*, *EYA1*, *EYA2*, *FAM49A*, *FOXN3*, *LMNA*, *MYO5A*, *PAX3*, *PCDH15*, *RTTN*, *SMAD1*, *XXYLT1* and *ZEB1*.**

**Figure S19. Pie chart, illustrating protein product classification of the human genes, harbouring genomic markers in significant association with craniofacial phenotypes. The genes are: *AGXT2*, *BMP4*, *CACNB4*, *COL11A1*, *EGFR*, *EYA1*, *EYA2*, *FAM49A*, *FOXN3*, *LMNA*, *MYO5A*, *PAX3*, *PCDH15*, *RTTN*, *SMAD1*, *XXYLT1 and ZEB1*.**

---

**Table S1.Manually annotated facial landmarks used in the study.**

**Table S2.Genetic associations with pigmentation traits.**

Gene: gene name; rs#: reference SNP ID number; SNP: chromosomal location of the marker; Genomic annotation: genomic location of the marker; UNADJ: Unadjusted p-values; BONF: Bonferroni single-step adjusted; HOLM: Holm (1979) step-down adjusted; SIDAK_SS: Sidak single-step adjusted; SIDAK_SD: Sidak step-down adjusted; FDR_BH: Benjamini & Hochberg (1995) step-up FDR control; FDR_BY: Benjamini & Yekutieli (2001) step-up FDR control.

**Table S3.Genetic syndromes displaying various craniofacial abnormalities, used to locate candidate genes for the study.**

## Appendix S1.

**Comprehensive list of web resources used for candidate gene search and its output. Note the presence of multiple tabs in this spreadsheet.**

## Appendix S2.

**Three spreadsheets, detailing a list of 8,518 genetic markers genotyped in 587 DNA samples and a list of 2,332 markers used for association analyses, following MAF (2%) and HWE filtering.**

